# Evolution of coprophagy and nutrient absorption in a cave salamander

**DOI:** 10.1101/123661

**Authors:** Daphne Soares, Rachel Roenfeldt, Shea Hammond, Michael E. Slay, Dante B. Fenolio, Matthew L. Niemiller

## Abstract

The transition from carnivory to omnivory is poorly understood. The ability to feed at more than one trophic level theoretically increases an animal’s fitness in a novel environment. Because of the absence of light and photosynthesis, most subterranean ecosystems are characterized by very few trophic levels, such that food scarcity is a challenge in many subterranean habitats. One strategy against starvation is to expand diet breadth. Grotto salamanders (*Eurycea spelaea*) are known to ingest bat guano deliberately, challenging the general understanding that salamanders are strictly carnivorous. Here we tested the hypothesis that grotto salamanders have broadened their diet related to cave adaptation and found that, although coprophagous behavior is present, salamanders are unable to acquire sufficient nutrition from bat guano alone. Our results suggest that the coprophagic behavior has emerged prior to physiological or gut biome adaptations.

## Introduction

Coprophagy is a feeding strategy commonly found in invertebrates (Weiss 2006), but much less so in vertebrates. Coprophagy sometimes exists in mammals such as rodents and lagomorphs, and to a lesser degree in pigs, horses, dogs and nonhuman primates. In amphibians, coprophagy is rare but when present may influence larval development of some species with herbivorous larvae. For example, herbivorous tadpoles regularly feed on feces of conspecifics in captivity (Gromko et al. 1973; Steinwascher 1978; Pryor and Bjorndal 2005), even when other food sources are available ad libitum (Pryor and Bjorndal 2005). Herbivorous tadpoles have digestive morphologies and physiologies similar to other herbivorous vertebrates that rely on hindgut fermentative digestion (Pryor and Bjorndal 2005) and ingest feces to inoculate their digestive tracts with beneficial microbes (Steinwascher 1978; Beebee 1991; Beebee & Wong 1992). Growth rates are slower when feces are removed from the diet (Steinwascher 1978), and feces are lower in energy content after reingestion (Gromko et al. 1973). These studies indicate that herbivorous tadpoles benefit nutritionally from coprophagy.

In predatory amphibians, coprophagy is exceedingly rare. However, faces consist of a readily available food resource for animals living in energy-limited environments, such as caves. Food and nutritional resources in caves are derived from surface inputs and can be limited both temporarily and spatially within these systems (Culver and Pipan 2014). Likewise, foraging in aphotic habitats of caves presents significant challenges for animals that potentially may go weeks to months between feeding bouts. Guano produced by seasonally roosting bats represents an important food source for both terrestrial and aquatic invertebrates (Howarth 1983; Poulson and Lavoie 2000), which in turn are prey for fishes and salamanders (Poulson and Lavoie 2000; Graening 2005; Niemiller and Poulson 2010; Fenolio et al. 2006, 2014). Salamanders have been known to be strictly carnivorous but Fenolio et al. (2006) showed that obligate cave-dwelling Grotto Salamander larvae (*Eurycea spelaea*, Figure 1) ingests bat guano. This behavior is not incidental to the capture of aquatic invertebrate prey. Stable isotope signatures suggest nutrients from bat guano could be incorporated into salamander tissues, and nutritional analyses revealed that bat guano is comparable to potential prey items in nutritional and energy content, suggesting that bat guano could be a viable alternative food source in some energy-poor cave systems. Since the relative importance of guano in the diet of subterranean salamanders is unknown, the aim of this study was to determine whether subterranean salamander larvae could persist on an exclusive guano diet compared to the typical carnivorous diet of salamanders.

**Figure 1.**
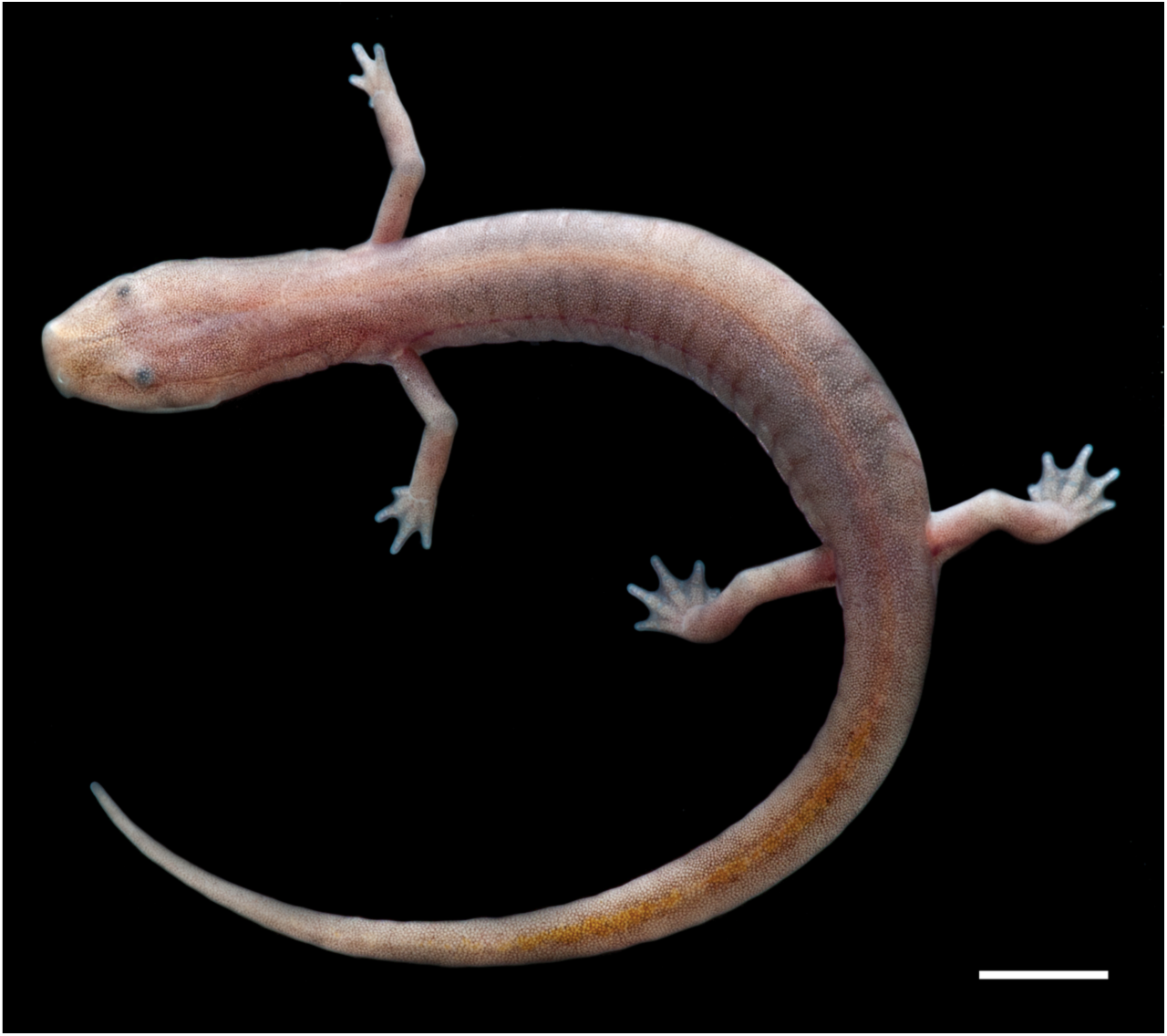
*Eurycea spelaea* showing troglobitic characters, lack of pigmentation and microphthalmy.Scale bar = 0.5cm

## Methods

All experiments were conducted under the approval of animal protocol #15022 by the Rutgers Newark Institutional Animal Care and Use Committee which handles NJIT research. We collected 46 Grotto Salamanders from January-Stansbury Cave located in the Ozark Plateau National Wildlife Refuge in Delaware County, Oklahoma. The cave contains a maternity colony (ca. 15,000 individuals) of federally endangered Gray bats (*Myotis grisescens*) from late April through early November (Fenolio et al. 2006, 2014). Salamanders were housed individually in mesocosms submerged in the cave stream and fed every four days either a strict diet of live amphipods, bat guano, or nothing for study duration. Amphipods and bat guano were collected fresh on the day of feeding from the cave. Salamanders were randomly assigned to one of two prey types and one of four feeding treatments based on percentage of initial body mass: 0% (control), 2.5%, 5%, and 10%. Salamanders were massed before feeding to track body mass loss or gain and fed the corresponding percentage of initial body mass of amphipods or guano. Care was taken to ensure that salamanders ate all food, and to remove any food remnants before the next feeding. Salamanders that lost ≥30% of initial body mass were removed the study. We used ANCOVA to compare body mass of the different treatments in MatLab. Salamanders were released back into the cave after the study per permitting regulations.

## Results

<u>Treatment groups:</u> All treatment groups lost some body mass during the study (34 days; Figure 2). Animals in the control group (n=10) were removed from the study earlier (27 days) than the other groups (34 days) due to body mass loss. Salamanders in the control group experienced the steepest loss of body mass. In general, salamanders fed guano lost more body mass than salamanders fed amphipods, and at 34 days, most guano-fed salamanders had reached the 30% loss limit. Body mass was more variable in amphipod-fed groups with both gains and losses. Salamanders fed 2.5% of initial body mass (IBM) lost 30.22% body mass eating guano compared to 10.35% eating amphipods. Salamanders fed 5% IBM lost 39.19% body mass when eating guano compared to 7.3% eating amphipods. Body mass loss was least for salamanders fed 10% IBM and guano-fed salamanders lost 27.79% body mass compared to 8.54% for the amphipod group. <u>Comparisons of weight loss:</u> For salamanders fed 2.5% IBM, body mass loss rates for guano-fed and amphipod-fed groups were slower than the control group (Guano-fed: F=6.82, p=0.01; Amphipod-fed: F=12.14, p=0.0007) but not different from each other (F=2.86, p=0.09). For salamanders fed 5% IBM, guano-fed animals lost body mass at a slower rate than amphipod-fed animals (F=11.05, p=0.0012) and control animals (F=14.75, p=0.0002), while amphipod-fed animals lost body mass similarly to control animals (F=0.07, p=0.795). For salamanders fed 10% IBM, the amphipod-fed group lost body mass slower than the guano-fed group (F=6.4, p=0.131) and control group (F=26.26, p=1.09e- 6), while the guano-fed group was similar to the control group (F=8.02, p=0.005).

**Figure 2.**
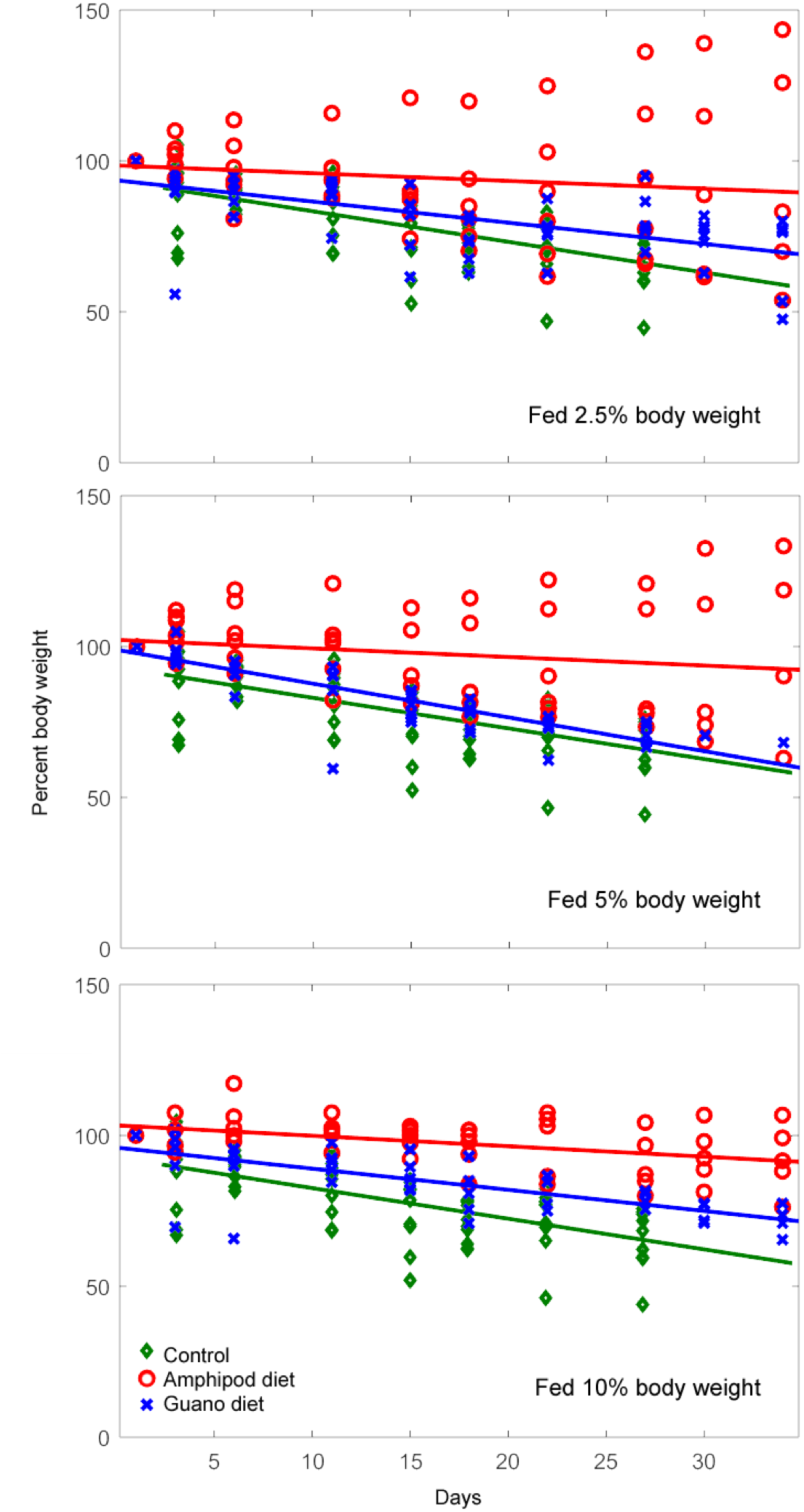
Regression lines based on body mass loss of different diet types and amounts.Salamanders were fed nothing (green), live amphipods (red) or guano (blue). Groups were fed every four days based on their initial body weight, with 2.5% (A), 5%(B) or 10% (C). The calculated regression lines were as follows: Control -1.16x+96.01 R^2^=0.54, n=10; 2.5%_amphipod_ - 0.26x+98.49, R^2^=0.39, n=6; 2.5%_guano_ -0.70x+93.58, R^2^= 0.02, n=6; 5%_amphipod_ -0.28x+102.22, R^2^=0.03, n=6; 5%_guano_ -1.12x+98.89, R^2^=0.77, n=6; 10%_amphipod_ -0.35x+103.36, R^2^=0.21, n=6; 10%_guano_ -0.70x+96.01, R^2^=0.53, n=6.

## Discussion

Shifts in habitat are often linked with dietery shifts, as environmental changes frequently cause organisms to alter foraging behaviors (Rosalino et al. 2005; McMeans et al 2015). The transition from surface to subterranean habitats involves dramatic morphological, physiological, and behavioral changes associated with life in complete darkness and often limited energy resources, including a predicted increase in dietary breadth (Culver 1982, 1994; Holyoak & Sachdev 1998; Fenolio et al. 2006). In subterranean salamanders, coprophagy may be an unusual foraging strategy to exploit a nutritious and seasonally abundant resource (i.e., bat guano) in an otherwise food-limited environment. While it has been demonstrated that Grotto salamander larvae will regularly employ coprophagy of calorically-rich bat guano (Fenolio et al. 2006), our study suggests that Grotto salamander larvae are unable to thrive on a guano-exclusive diet for a prolonged period. Ingesting greater amounts of guano slowed rates of body mass loss, but all salamanders within our three treatments (2.5%, 5%, and 10%) lost 30% of initial body mass within 34 days.

The disconnect between coprophagous behavior in Grotto salamanders and the lack of apparent absorption may have several possible explanations. First, Grotto salamander larvae, and salamanders in general, do not possess the morphological and physiological digestive traits necessary to exploit guano as a food resource. Salamanders in general are strict carnivores with short digestive tracts and have buccal enzymes with low amylolytic activity (Stevens and Hume 2004). In contrast, coprophagy is most often associated with herbivory, which predominately utilize post-gastric (hindgut) fermentation and the consumption of feces increases the absorption of nutrients and inoculate the hind gut with microbes (Claus et al. 2007). The selective consumption of predigested material is a form of omnivory. We know relatively little about the adaptive advantages of and the selective drivers that favor omnivory, and by proxy coprophagy, in vertebrates (but see Diehl 2003). Coprophagy requires the evolution of not only a coprophagous behavior but also the evolution of morphological and physiological digestive traits to process feces. It is unknown whether these traits are linked, but theoretically behavioral evolution can precede physiological and morphological evolution. Second, since Grotto salamanders are ingesting feces with high protein content (54%; Fenolio et al. 2006) of insectivores (bats) rather than feces from herbivores, a vastly different gut microbiome is needed to efficiently digest feces. So in addition to lacking the morphological and physiological traits, Grotto salamanders may not possess the necessary gut flora to digest and fully process the contents of bat guano. Ley et. al (2009) found that diet can impact gut microbiome diversity in mammals, which increases with evolution from carnivory to omnivory. Digestive evolution in amphibians, as well as their gut biomes and the gut’s propensity for evolution, is yet to be examined in detail. Finally, coprophagy may reflect mistaken identity due to an innate feeding response for moving prey. In subterranean habitats, aquatic salamanders and cavefishes rely heavily on mechanosensation to detect and capture moving prey. Guano falling into a pool and settling on the substrate may elicit a similar feeding response as crustaceans and other aquatic invertebrates. Guano may not be immediately rejected but ingested instead because of the high protein and fat content of the insectivorous guano. Alternatively, guano may possess a micronutrient, vitamin or mineral otherwise scarce in the subterranean habitat (see Fenolio et al. 2006). While guano may not prevent a loss in mass, it may still offer some nutritional benefit.

## Acknowledgements

We thank Dr. Gal Haspel for Matlab scripts and comments, Ozarks Plateau National Wildlife Refuge for lodging and cave access, Sheilah Roenfeldt for field assistance, and Oklahoma Department of Wildlife Conservation for collection and research permits.

## Author contributions

DS was responsible for conception, design, interpretation and drafting; RR, SH and MES were responsible for the execution and drafting; MLN was responsible for interpretation and drafting; DF was responsible for conception, design and drafting of the work.

## Data availability

All data associated with this study are available from the figshare digital repository: 10.6084/m9.figshare.4805656

